# The molecular mechanism of load adaptation by branched actin networks

**DOI:** 10.1101/2021.05.24.445507

**Authors:** Tai-De Li, Peter Bieling, Julian Weichsel, R. Dyche Mullins, Daniel A. Fletcher

**Affiliations:** Department of Bioengineering & Biophysics Program, University of California, Berkeley, CA 94720, USA; Division of Biological Systems & Engineering, Lawrence Berkeley National Laboratory, Berkeley, CA 94720, USA; Advanced Science Research Center, City University of New York, New York, NY, USA; Department of Cellular and Molecular Pharmacology and Howard Hughes Medical Institute, University of California, San Francisco, CA 94158, USA; Department of Systemic Cell Biology, Max Planck Institute of Molecular Physiology, Dortmund, Germany; Department of Chemistry, University of California, Berkeley, 94720, USA; Chan Zuckerberg Biohub, San Francisco, CA 94158, USA

## Abstract

Branched actin networks are self-assembling molecular motors that move biological membranes and drive many important cellular processes. Load forces slow the growth and increase the density of these networks, but the molecular mechanisms governing this force response are not well understood. Here we use single-molecule imaging and AFM cantilever deflection to measure how applied forces affect each step in branched actin network assembly. Unexpectedly, force slows the rate of filament nucleation by promoting the interaction of nucleation promoting factors with actin filament ends, limiting branch formation. This inhibition is countered by an even larger force-induced drop in the rate of filament capping, resulting in a shift in the balance between nucleation and capping that increases network density. Remarkably, the force dependence of capping is identical to that of filament elongation because they require the same size gap to appear between the filament and load for insertion. These results provide direct evidence that Brownian Ratchets generate force and govern the load adaptation of branched actin networks.

## Introduction

Unlike conventional motor proteins, which produce force by converting chemical energy into conformational changes, branched actin networks generate force by spatially and temporally coordinating interactions between a set of four core components: (i) a nucleation promoting factor (NPF) related to the Wiskott-Aldrich Syndrome Protein (WASP), (ii) the Arp2/3 complex, (iii) capping protein (CP), and (iv) profilin-actin complexes (Achard et al., 2010; Akin and Mullins, 2008; Bieling et al., 2018; Loisel et al., 1999). Branched actin network assembly begins when signaling molecules, such as Rho-family GTPases, cluster together and activate NPFs on a membrane surface (Dominguez, 2009; Husson et al., 2010). Active NPFs locally promote actin nucleation and branching by the Arp2/3 complex (Mullins et al., 1998; Rohatgi et al., 1999). These newly created actin filaments elongate at their fast-growing (barbed) ends from profilin-actin complexes (Funk et al., 2019), fed by an intrinsic polymerase activity of the nucleation promoting factors (Bieling et al., 2018). Individual filaments elongate and push against the NPF-coated membrane surface, but only for a short time before capping protein terminates their growth (Edwards et al., 2014; Schafer et al., 1996). As a result, steady-state network growth requires continual nucleation. Whenever active NPF molecules are concentrated together on a membrane surface, this sequence of interactions is sufficient to create a powerful molecular motor capable of generating kilopascal (nN/*μ*m^2^) pressures (Bieling et al., 2016; Marcy et al., 2004; Parekh et al., 2005).

The response of a molecular motor to force is fundamental to understanding its biophysical mechanism. Force, for example, coordinates the out-of-phase stepping of two-headed kinesins along a microtubule (Yildiz et al., 2008), and it can cause dynein motors to step ‘backward,’ toward the microtubule plus end (Gennerich et al., 2007). Force also causes some myosin motors to cling more tightly to actin filaments (Laakso et al., 2008). Similarly, force produces dramatic effects on the motor activity of branched actin networks, increasing the number and density of actin filaments (Bieling et al., 2016; Mueller et al., 2017) as well as the mechanical efficiency of the network (Bieling et al., 2016). These changes in filament density also optimize the material properties of a branched actin network to better resist deformation and to generate higher forces. Load adaptation in a growing actin network comprises two distinct processes: (i) reorientation of filaments within the network (Mueller et al., 2017; Weichsel and Schwarz, 2010) and (ii) an increase in the steady-state number of growing filaments (Bieling et al., 2016). Filament reorientation has been explained by kinetic competition models (Weichsel and Schwarz, 2010) but the mechanism underlying the force-induced increase in filament number remains unknown.

To figure out how load forces increase the number of growing filaments, we studied branched actin networks assembled from purified proteins on micro-patterned, functionalized glass surfaces. We measured the rate of network growth against defined forces using a modified Atomic Force Microscope (AFM) and simultaneously visualized the flux of constituent molecules -actin, capping protein, and the Arp2/3 complex- into the network by total internal reflection fluorescence (TIRF) microscopy (Bieling et al., 2016). From these measurements, we find that load adaptation in the network arises from a mismatch in the force-dependent activities of capping protein and the Arp2/3 complex. Contrary to our expectations, new branch formation is reduced by load-dependent interaction with nucleation promoting factors on the membrane, a process we refer to as ‘barbed end interference’. However, rates of capping are reduced even more significantly, leading to a net increase in actin density with increasing force. Interestingly, we show that the force responses of filament elongation and capping are closely matched, ensuring that the average filament length and molecular stoichiometry of the network remain constant across a wide range of load forces. The exponential force response of filament capping suggests that it is governed by a Brownian Ratchet similar to the one that is thought to govern filament elongation. To test this idea directly we created a ‘bulky’ capping protein mutant that adds a larger length increment to the barbed end of an actin filament. When added to freely growing actin filaments in solution, this bulky mutant shows no defects in capping activity, but when added to actin networks growing against a load force it exhibits a dramatically different force response and cannot keep pace with wildtype capping protein. Overall, our work reveals that a combination of matched and mismatched force responses work together to stabilize branched actin networks and enable them to respond to changing load forces.

## Results

### Load forces produce different effects on actin filament nucleation and capping

The steady-state density of free barbed ends (*μ*m^−2^) at the growing surface of a branched actin network is determined by the ratio of the nucleation rate across that surface (*μ*m^−2^sec^−1^) divided by the per-filament capping rate (sec^−1^; (Mullins et al., 2018)). To change the density of growing filament ends, load forces must somehow alter the balance between nucleation and capping. Some numerical simulations propose that the rate of filament nucleation increases under load as a result of the autocatalytic nature of Arp2/3 branching (Carlsson, 2003), but this has never been tested experimentally. Furthermore, as far as we can tell, the force-dependence of filament capping has never been investigated or incorporated into numerical simulations.

We measured the rates of filament nucleation, elongation, and capping in branched networks under load by performing TIRF microscopy on fluorescently labeled Arp2/3 complexes, actin, and capping protein (Figure 1A, B) as these molecules incorporated into networks growing from coverslips printed with square patterns of the C-terminal, Arp2/3-activating region of the nucleation promoting factor WAVE1 (WAVE1ΔN). We applied defined load forces using a calibrated AFM cantilever and found that the density of all three network components increased monotonically with increasing loads (Figure 1C). We calculated the rate of Arp2/3 complex incorporation by multiplying the average fluorescence intensity in the TIRF field (Figure 1B) by the steady-state network growth velocity under each loading condition (Figure 1C). Contrary to previous predictions, the rate of Arp2/3 incorporation decreased linearly with applied force (Figure 1D), falling to approximately 50% of its initial value at high load (~1200 Pa).

**Figure 1.**
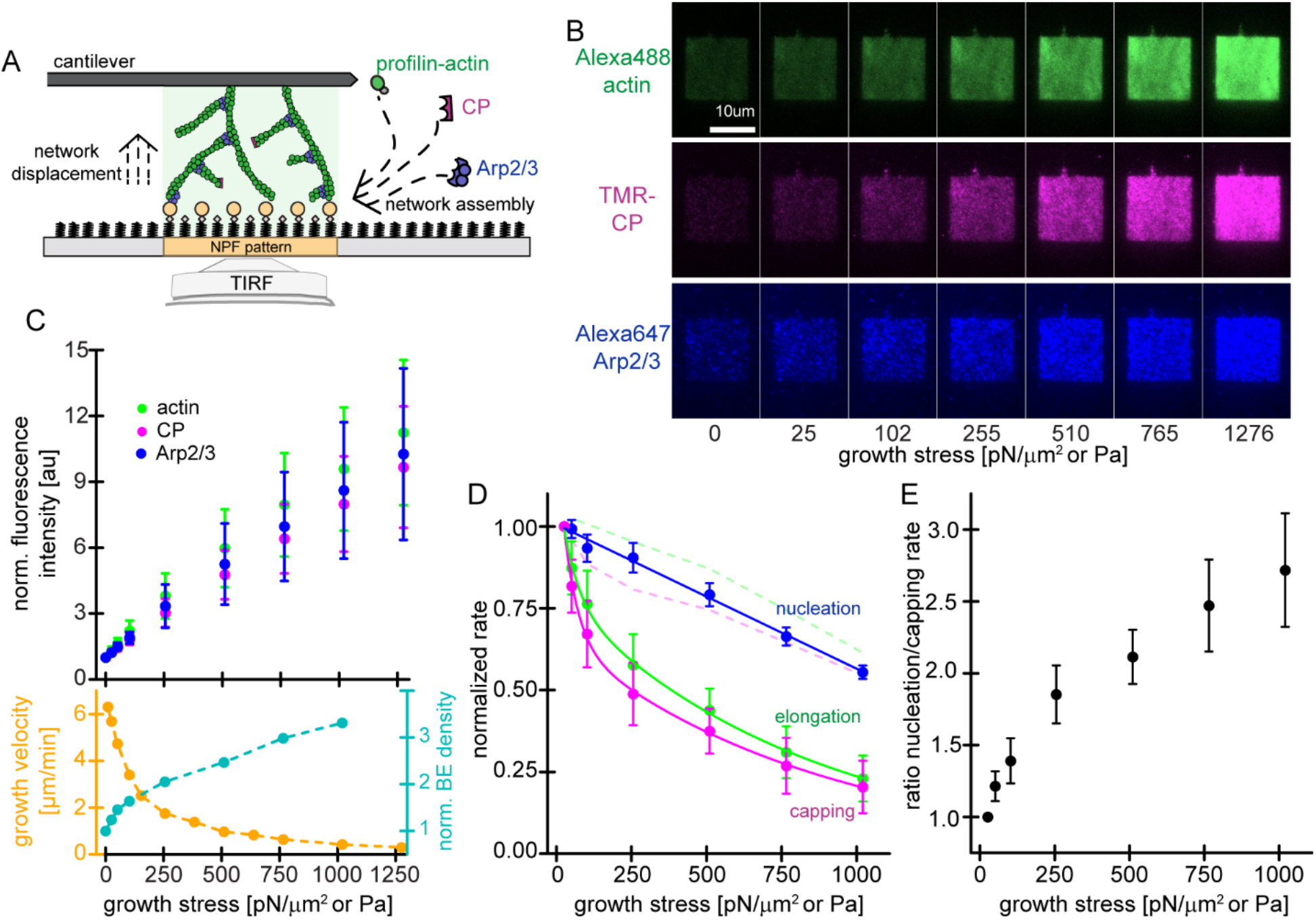
Effect of mechanical load on Arp2/3-dependent nucleation. **A)** Schematic illustration of actin networks generated by profilin-actin, the Arp2/3 complex and capping protein from surfaces coated with NPF (WAVE1ΔN). Conditions are 5μM actin (1% Alexa 488-labeled), 5μM profilin, 100nM Arp2/3 (5% Alexa647-labeled), 100nM CP (15% TMR-labeled) if not indicated otherwise. **B)** TIRFM images of Alexa488-actin (top), TMR-CP (middle) and Alexa647-Arp2/3 (bottom) incorporation into dendritic actin networks at indicated growth stress. **C)** Top: Quantification of florescent intensities for indicated network components as a function of applied load. Bottom: Corresponding growth velocities and free barbed end densities as measured in (Bieling et al., 2016). Error bars are SD. **D)** Average rates of filament elongation, capping and nucleation (calculated by the product of the bulk fluorescence intensities and the network growth velocity, normalized to the flux at 25pN/μm2) as a function of external load. Capping and elongation rates were obtained by normalizing their network incorporation rates (dashed lines) by the relative density of free barbed ends as shown in C). Error bars are SEM. **E)** Ratio of nucleation and capping rates as a function of external load. Error bars are SEM.

To quantify the effect of force on filament elongation and capping, we also multiplied the average fluorescence intensities of labeled actin and capping protein by the network growth rate. Because both actin and capping protein interact with free filament ends whose number increases with load force, we also normalized the bulk incorporation rates by the free barbed ends densities produced under the same loads (Figure 1C; (Bieling et al., 2016)). This analysis revealed a dramatic, force-induced reduction in the rates of filament elongation and capping (Figure1 D), which can be fit by a double exponential decay. When load force is converted to force per filament, however, the responses of both elongation and capping are well fit by a single exponential decay (Supplemental Figure 1), suggesting that the processes are governed by the same mechanism. Importantly, the drop in filament capping outstrips the drop in nucleation rate, causing the nucleation/capping ratio to increase with increasing force (Figure 1E). We conclude, therefore, that the force-induced increase in filament number is driven by the mismatched force responses of filament nucleation and capping rates, not by a force-dependent increase in the rate of nucleation.

Our measurements are not explained by a previous model for the effect of force on actin nucleation (Carlsson, 2003), so we investigated the molecular mechanism responsible for this effect. We first visualized incorporation of individual Arp2/3 complexes into growing networks (Figure 2A) using TIRF microscopy. For these single-molecule experiments, we mixed trace amounts of fluorescent Arp2/3 with a large excess of unlabeled complexes (1:5000) and used this mixture to form branched actin networks from WAVE1ΔN-coated surfaces. At this low labeling ratio, individual fluorescent Arp2/3 complexes appeared abruptly as diffraction-limited spots on the WAVE1ΔN-coated surface and then faded as they moved with the growing actin network away from the coverslip surface and out of the TIRF illumination field (Figure 2A, Supplemental Movie 1). We classified spots whose intensity decayed exponentially with time as network incorporation events, and we rejected fluorescent spots that disappeared in a single step due either to dissociation or photobleaching. Based on the measured surface density of NPFs in our assay (1850 um^−2^, (Bieling et al., 2018)), we determined a nucleation rate of 0.037 s^−1^ per WAVE1ΔN molecule. This rate is notably faster than the limiting step in Arp2/3 branching observed in solution assays (Helgeson and Nolen, 2013; Smith et al., 2013; Zalevsky et al., 2001), demonstrating that nucleation within a branched network is surprisingly rapid. However, because release of the NPF from the nascent branch appears to limit nucleation in solution (Helgeson and Nolen, 2013; Smith et al., 2013), we speculate that retrograde forces generated by network growth facilitate dissociation of the surface-bound NPF and accelerate nucleation in the context of a force-generating network.

**Figure 2.**
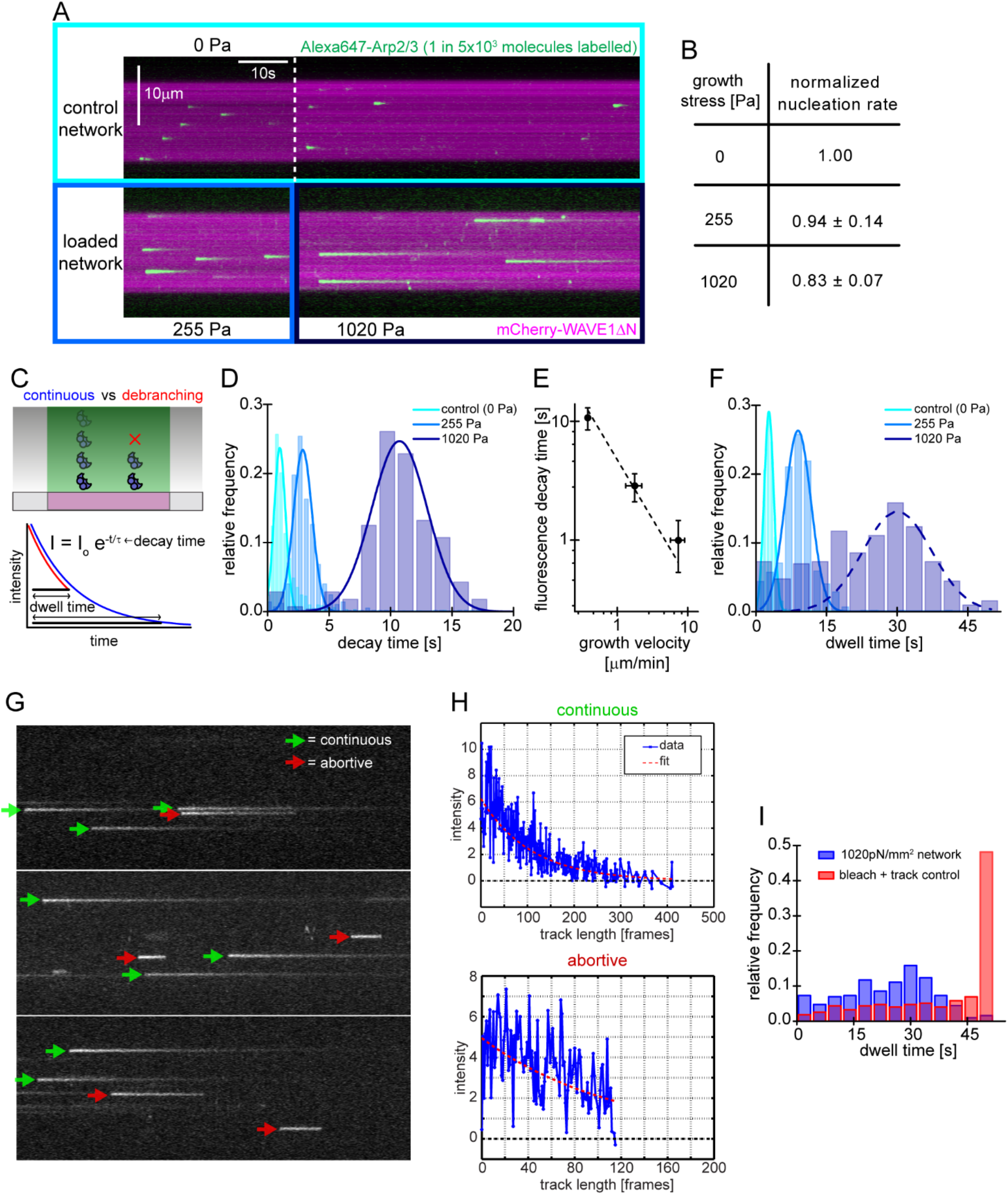
Single molecule characterization of force-dependent Arp2/3 nucleation. **A)** Kymographs of single molecule nucleation on NPF surfaces (mCherry-WAVE1ΔN, magenta) by spike-in of a small fraction of Alexa647-Arp2/3 (green, c=20 pM) into the overall Arp2/3 pool (100nM) at indicated applied stress (lower panel) or in an adjacent unloaded control network (upper panel). **B)** Mean nucleation rate by single molecule imaging normalized to the nucleation rate in an adjacent unloaded control network at indicated growth stress. Error bars are SEM. **C)** Scheme of single molecule Arp2/3 nucleation by TIRFM for either continuous (blue) or debranching (red) scenarios. Individual intensity trajectories are characterized by either their transit time through the evanescent field or their actual dwell time. The latter is sensitive to debranching. **D)** Normalized frequency of fluorescence transit times of single Alexa 647-Arp2/3 molecules in networks assembled at indicated stress. Continuous lines are Gaussian fits to the data. **E)** Double-logarithmic plot of the mean fluorescence transit time (+/− SD) as a function of network growth velocity. The dashed line show perfect reciprocal correlation (slope = −1). **F)** Normalized frequency of fluorescence dwell times of single Alexa 647-Arp2/3 molecules in networks assembled at indicated stress. Continuous lines are Gaussian fits to the data. Dashed line is a Gaussian fit to a partial (normally distributed) fraction of the 1020 pN/μm2 data set. **G)** Examples of kymographs from TIRF microscopy of individual Arp2/3 molecules in networks under high load (1020pN/m2). Individual complexes are either continuously moving towards the rear of the evanescent field (continuous, green arrows) or dissociating prematurely (abortive, red arrows). **H)** Representative time courses of fluorescence intensity for individual Arp2/3 complexes as a function of number of imaging frames categorized as either continuous (top panel) or abortive (bottom panel). **I)** Relative frequency of dwell times for Arp2/3 molecules in dendritic networks at high load (1020pN/m2, blue) compared to the bleaching and loss of tracking control (red, see Supplementary Figure 1). Note that the frequency of early loss events is exceeding the combined bleaching and tracking loss frequency.

Increasing load forces caused the fluorescence decay of individual Arp2/3 molecules to slow down, as expected from load-dependent slowing of network movement (Figure 2A, Supplemental Movie 1, (Bieling et al., 2016)). We calculated a time constant for each Arp2/3 incorporation event by fitting its fluorescence decay profile with a single exponential (Figure 2C). The calculated time constants for each loading condition followed a Gaussian distribution (Figure 2D) whose mean was inversely correlated with network growth velocity (Figure 2E). In these single-molecule measurements, the rate of Arp2/3 complex incorporation decreased moderately (~17%) with load (Figure 2B).

Although both single-molecule and bulk fluorescence measurements revealed a decrease in the rate of Arp2/3 incorporation under load (Figure 1D, 2B), the decrease determined by bulk fluorescence (~50%) was more pronounced than for single molecules (~17%) over the same stress range (0 to 1000 Pa). To see whether this difference reflects rapid dissociation of a fraction of Arp2/3 complexes from the network, we measured the dwell time between appearance and disappearance of each molecule (Figure 2F, Supplemental Methods). Under low and intermediate loads (25 and 255 Pa), the distribution of Arp2/3 dwell times is approximately Gaussian, and the mean lifetime increases with decreasing network velocity (Figure 2F). At high loads (1020 Pa), however, the distribution develops a ‘shoulder’ of dwell times shorter than expected (Figure 2F). Fluorescence decay curves of these short-lived events resemble truncated exponentials, consistent with abrupt dissociation of the Arp2/3 complex (Figure 2G,H). These events were too frequent to be accounted for by experimental artifacts such as photobleaching and tracking glitches (Figure 2I, Supplemental Figure 2). Taken together, these results reveal that a significant fraction of Arp2/3 complexes dissociate from the network under high load. Whether forces increase the rate of Arp2/3 dissociation as indicated by recent experiments pulling on individual branches (Pandit et al., 2020) or whether the slower rate of network growth under stress simply keeps the ephemeral branches within the evanescent field for a longer time remains to be determined.

### Effect of load on actin-WH2 interactions

Why does compressive loading reduce the rate of Arp2/3-dependent nucleation? In addition to promoting nucleation by delivering actin monomers to the Arp2/3 complex, WASP-family proteins associate with actin filament barbed ends via their WH2 domain and thus tether dendritic networks to the boundary they push against (Co et al., 2007; Funk et al., 2021). This means that free barbed ends and actin monomers are in direct competition to occupy available NPF WH2 domains (Figure 3A).

**Figure 3.**
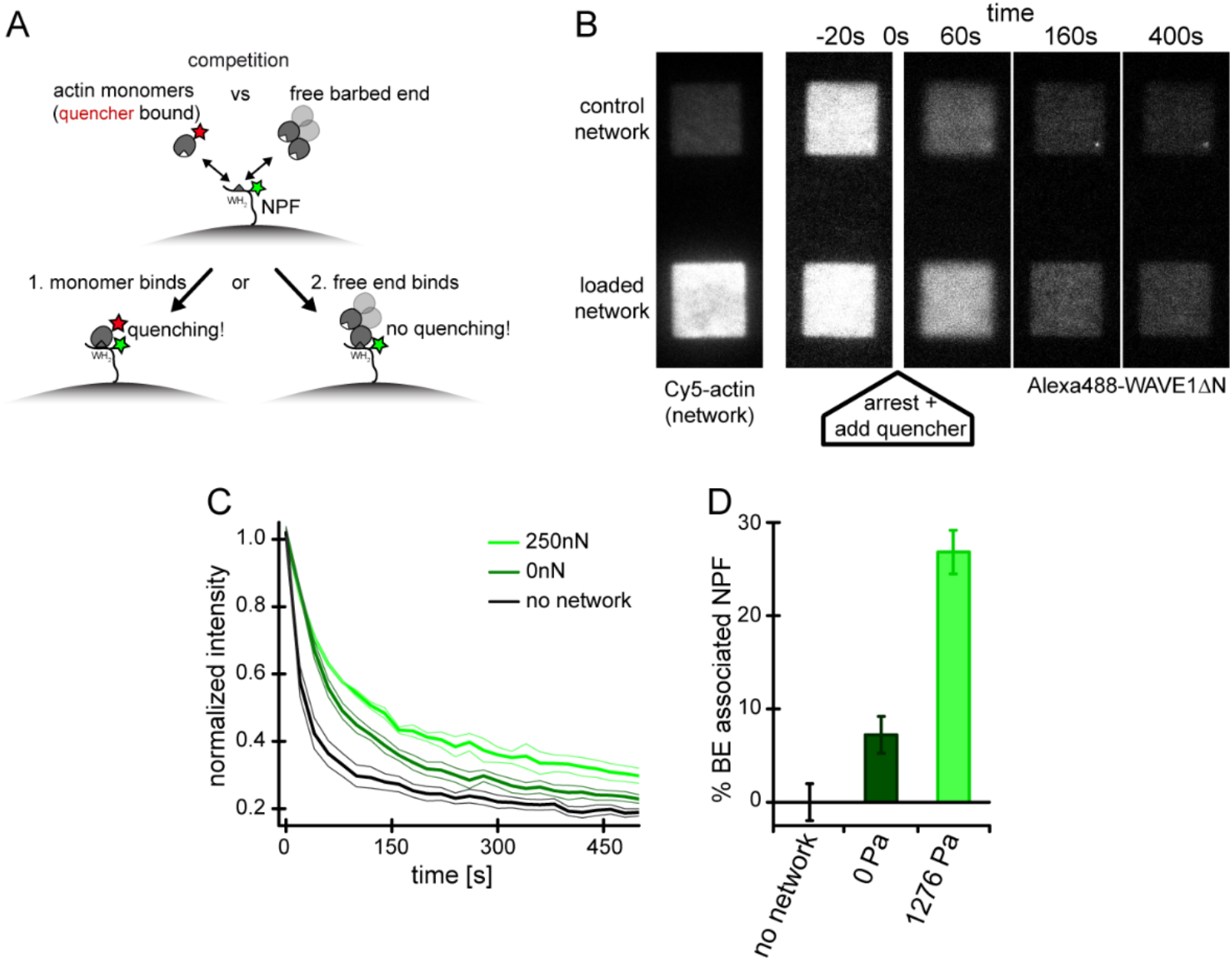
Free barbed ends attach to and sequester the WH2 domain of the NPF in a load-dependent manner. **A)** Scheme of the FRET setup. Surface-bound, donor- (Alexa488-) labeled NPF molecules can interact with either quencher- (Atto540Q-) labeled actin monomers resulting in decrease of donor fluorescence or unlabeled terminal protomers of uncapped barbed ends resulting in no change in fluorescence. The terminal protomers are unlabeled since quencher-labeled monomers are introduced only upon network arrest. **B)** Time lapse TIRF microscopy images of Alexa 647-Actin (left image) or Alexa 488-WAVEΔN (FRET donor, other images) at indicated times after addition of 200μl fixation and quenching mix (t=0, 30 μM LatB, 30μM phalloidin, 5μM Atto540Q-actin, 7μM profilin, 37.5 μM myotrophin/V1 (CP inhibitor)) to 100 μl network assembly mix. **C)** Time-courses of the Alexa 488-WAVEΔN signal following the addition of quencher-labelled monomers at t=0 as shown in B) Error indicators are SD. **D)** Mean fraction of barbed end-associated NPF molecules in either in the absence of an actin network (black) or in the presence of a non-loaded (dark green) or 200nN loaded (light green) network (See Supplemental Methods). Error bars are SEM.

Because the concentration of surface-proximal free barbed ends rises in a load-dependent manner (Figure 1C, (Bieling et al., 2016)), we asked whether this might reduce the availability of the NPF for monomer binding. We adapted a recently developed Förster resonance energy transfer (FRET) assay (Bieling et al., 2018) to directly measure partitioning of WH2 domains between actin monomers and filament ends (Figure 3A). Briefly, we labeled a WH2-adjacent site of WAVE1ΔN with a fluorescent donor (Alexa488) and conjugated a non-fluorescent acceptor (Atto540Q) to monomeric actin using a labeling protocol that does not perturb binding to profilin and WH2 domains (Bieling et al., 2018; Funk et al., 2021).

To measure partitioning of WH2 domains between actin monomers and filament ends we micro-patterned glass coverslips with WAVE1ΔN, doped with 30% donor-labeled molecules. We then assembled dendritic networks under zero or high load (1276 Pa) in the absence of quencher-labeled actin (Figure 3B). We simultaneously arrested network growth and inhibited filament capping by adding Latrunculin B and Phalloidin together with myotrophin/V-1, a combination of inhibitors that arrests network growth but preserves free barbed ends (Bieling et al., 2016). At the same time, we also added quencher-labeled actin monomers, bound to Latrunculin B to prevent their incorporation into free filament ends. The quencher-labeled monomers induced a rapid drop in donor fluorescence (Figure 3B, Supplemental Movie 2) as they bound available WH2 domains. We interpret donor fluorescence remaining at long time scales to reflect WH2 domains protected from quencher-labeled monomers by interaction with filament barbed ends.

To quantify the fraction of WH2 domains bound to free barbed ends, we compared the remaining donor fluorescence of WAVE1ΔN micro-patterns in either the absence or presence of dendritic actin networks (Figure 3C). In unloaded networks, approximately 7% of WH2 domains are protected from quenching, while under high load the protected fraction increases by ~3.7-fold, to 27% (Figure 3D). These numbers are in striking agreement with the 3.3-fold increase in free barbed end density of networks under load (Figure 1C, (Bieling et al., 2016)). The 20% decrease in available WH2 domains also agrees well with the load-induced 17% drop in nucleation rate (Figure 2B). In summary, these results verify that applied forces raise the number of free barbed ends that in turn engage an increasing number of nucleation promoting factors in non-nucleating complexes (Funk et al., 2021; Mullins et al., 2018). This type of negative feedback mechanism, which we call ‘barbed end interference’ (Funk et al., 2021), quantitatively explains the force-induced decrease in Arp2/3-dependent nucleation.

### Filament capping follows the same force response curve as elongation

When comparing the effects of force on filament elongation and capping (Figure 1D), we noticed an almost identical force dependence, which explains our previous observation that the average filament length in a branched actin network does not change with applied load (Bieling et al., 2016). We confirmed that the average filament length in a growing network decreased as expected with increasing capping protein concentrations (Supplementary Figure 2). Elevated capping protein levels resulted in sparser networks of shorter filaments that grew faster at comparable loads (Supplementary Figure 2). In line with our previous work (Akin and Mullins, 2008), we observed that the increase in capping was entirely compensated for by elevated nucleation rates, confirming that capping protein stimulates nucleation in branched networks (Supplementary Figure 2, (Akin and Mullins, 2008; Funk et al., 2021)). Remarkably however, filament length remained force insensitive at all capping protein concentrations tested (Figure 4A).

**Figure 4.**
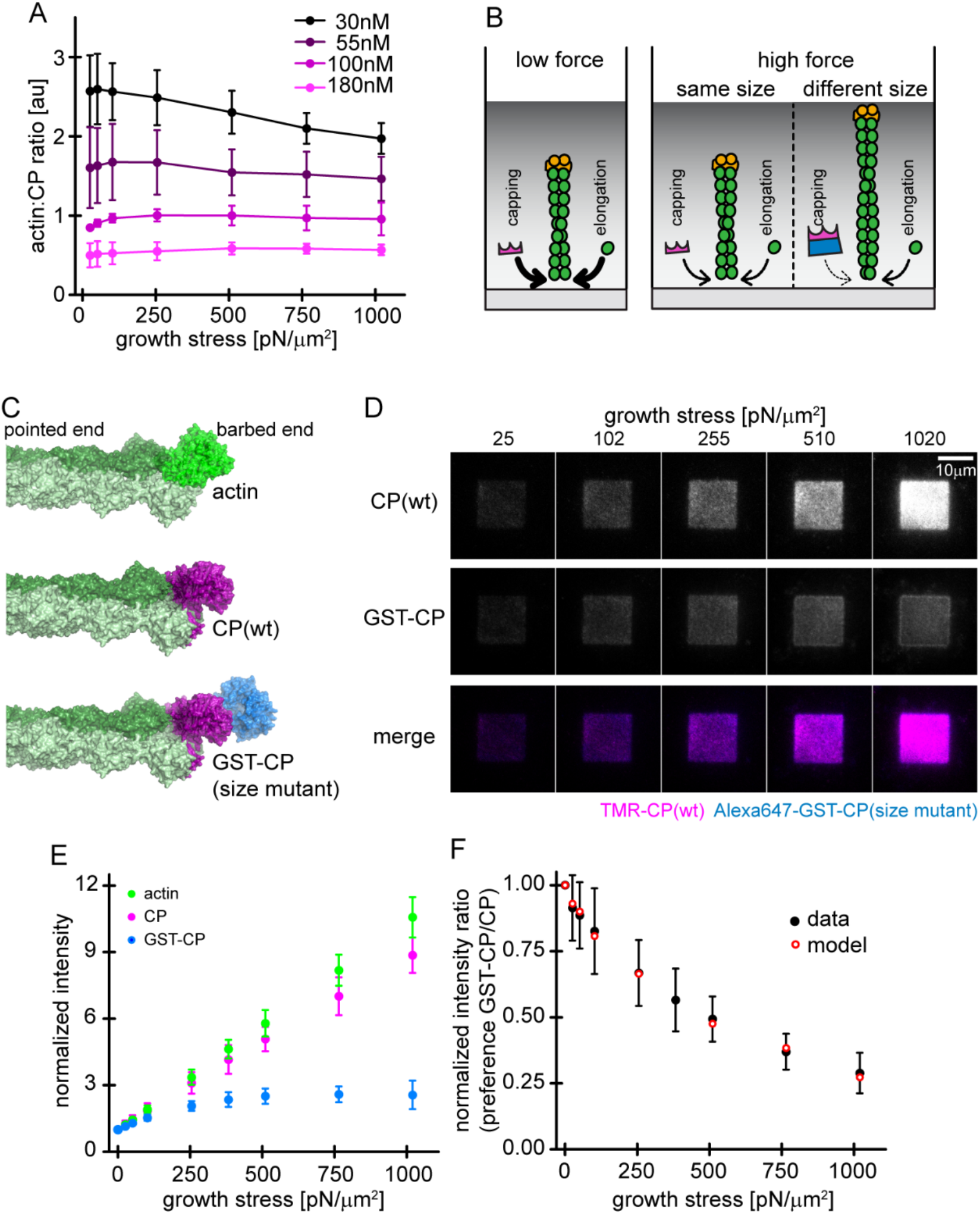
Load dependence of capping and a direct test of the elastic Brownian theory of force generation by actin networks. **A)** Ratios of capping protein to actin fluorescence for networks grown at different CP concentrations as indicated as a function of load (see Supplemental Figure 3). Error bars are SD. **B)** Illustration of the consequences of load dependence of capping and polymerization. Low load allows for high capping and polymerization rates (left panel). A similar load dependence of these two processes maintains filament length at high load (middle panel), whereas a difference in load dependence leads to changes in filament length (right panel). **C)** Structural models of a filament barbed end (light and dark green) bound by either an additional actin monomer (top panel, bright green), a wt CP heterodimer (middle panel, magenta) or an engineered GST and CP dimer fusion (“bulky variant”, bottom panel, magenta=CP, blue=GST). **D)** TIRFM images of dendritic actin networks (top panel=TMR-CP (wt), middle panel=Alexa647-GST-CP (bulky variant) and bottom panel=color merge) at indicated stress. Networks were assembled at standard conditions, except that CP (wt) concentration was 90nM (of which 10nM were TMR-CP) and Alexa647-GST-CP concentration was 10nM. **E)** Mean Alexa 488-actin, TMR-CP(wt) or Alexa 647-GST-CP (bulky variant) intensity normalized to the intensity of an adjacent unloaded network as a function of load. Error bars are SD. **F)** Measured mean fluorescence intensity ratios of CP(wt)/GST-CP(bulky variant) normalized to the intensity ratio of an adjacent unloaded network as a function of load. Error bars are SD. Red open circles are derived from the Brownian Ratchet Model (see Appendix).

Although not explicitly predicted, a link between filament capping and elongation is implicit in Elastic Brownian Ratchet theories of force generation (Mogilner and Oster, 1996; Peskin et al., 1993). According to these theories, the rate at which a protein binds the end of a filament growing against a boundary depends on how often thermal motion opens a large enough gap to accommodate the incoming molecule (Figure 4B). Intriguingly, based on their structures (Funk et al., 2021; Kim et al., 2010; Narita et al., 2006), it appears that an actin monomer and a capping protein heterodimer (Figure 4D, top and middle) both require nearly the same gap size to bind the barbed end of an actin filament. Altering the size of capping protein should, therefore, change the force response of filament capping.

To test this prediction we constructed a ‘bulky’ capping protein by fusing dimeric glutathione S-transferase (GST) to the C-termini of both α and β subunits of capping protein (Figure 4C, bottom). From existing structures, we predict that the gap required for this bulky variant to bind to the filament end should be significantly larger than for wildtype capping protein (Funk et al., 2021; Kim et al., 2010; Narita et al., 2006). Under load, these larger gaps open much less frequently, and so the rate of capping by the bulky variant should decrease much more strongly with applied force. We confirmed that the bulky capping protein caps filament barbed ends with wildtype kinetics in solution (Supplemental Figure 3). To directly compare the force sensitivities of wildtype and bulky capping protein under the same conditions, we constructed dendritic actin networks using mostly (90%) unlabeled wildtype capping protein, doped with small amounts (5% each) of wildtype and bulky capping proteins labeled with different fluorescent dyes. We labeled wildtype capping protein with tetramethyl-Rhodamine (TMR) and the bulky variant with Alexa-647. Under low load forces both wildtype and bulky capping proteins incorporated into the network, but only wildtype capping protein maintained a constant stoichiometry with actin under high load (Figure 4D). The bulky capping protein mutant was effectively excluded from the network by applied force (Figure 4D-F, Supplemental Movie 3). We tested for tag-specific effects by using a bulky NusA-tag instead of GST-GST at the N-terminus of capping protein, which resulted in a similar force-dependent reduction of incorporation (Supplemental Figure 3). To test whether these results can quantitatively be described by a Brownian Ratchet mechanism, we developed an analytic model of actin network growth under load (see Appendix). Briefly, we assumed that the capping rate declines exponentially with opposing force according to the Brownian Ratchet model with 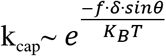, where the *K_B_* is the Boltzmann constant, *T* is the temperature, *θ* is the contact angle of filaments to the encounter surface, *f* is the force of one individual filament against the surface, and δ is the gap size required for incorporation of capping protein. The latter parameter differs for wildtype capping protein and its bulky variant, leading to their differential response to load. The average filament contact angle changes in response to force (Bieling et al., 2016; Mueller et al., 2017; Weichsel and Schwarz, 2010) and was estimated from the actin density and number of free barbed ends at various forces as measured previously (Bieling et al., 2016). To obtain the force *f* that opposes a single polymerizing filament, we divided the total external force on the whole network by the measured force-dependent number of free barbed ends sharing this load (Figure 1C; Supplemental Figure 1; (Bieling et al., 2016)). Finally, we took into account the internal tethering force that arises from the interaction between free barbed ends and NPF proteins on the surface, which in addition to the external load resists network movement (Figure 3). Except for this characteristic tethering force, all parameters are derived from experimental data. Remarkably, even with all but this single free parameter constrained, this model accurately matches the force- dependent change in the relative incorporation of wildtype capping proteins over its bulky variant (Figure 4E).

Furthermore, the characteristic tethering force yielded from this model (0.3 pN) falls within the typical range for weak protein-protein interactions. This data directly demonstrates that the addition of capping protein to the free barbed end of actin filaments in a branched network is size-dependent insertional process, whose force-dependent kinetics are described by Elastic Brownian Ratchet models (Mogilner and Oster, 2003, 1996)

## Discussion

### How free barbed ends interfere with activation of the Arp2/3 complex

One of the most unexpected results of our study is that force actually *decreases* Arp2/3 complex activity, suggesting that mother filament availability is not a limiting resource for the nucleation reaction. We show that the increased number of free barbed ends generated under load inhibits Arp2/3 complex activity by tying up WH2 domains and decreasing the ability of the NPFs to activate the Arp2/3 complex. Unlike the previously described ‘monomer gating’ mechanism (Akin and Mullins, 2008), in which free barbed ends compete with NPFs for monomeric actin, ‘barbed end interference’ is based on a direct interaction between NPFs and barbed ends of actin filaments (Funk et al., 2021; Mullins et al., 2018). This sort of negative feedback, in which the products of the nucleation reaction directly inhibit branching, also explains how Arp2/3-dependent nucleation achieves a constant, homeostatic rate despite its autocatalytic nature (Mullins et al., 2018).

### How force increases the number of free barbed ends

The force dependent increase in free barbed ends results from a change in the balance between nucleation and capping, driven by a large decrease in capping rate rather than by an increase in nucleation rate. Although the force sensitivity of filament capping has been neglected in most theoretical models of actin network assembly (Mogilner and Oster, 2003; Schaus et al., 2007), we can explain it using Elastic Brownian Ratchet theories developed to describe the effects of force on actin subunit addition during filament elongation (Mogilner and Oster, 1996; Peskin et al., 1993; Theriot, 2000).

Since both capping protein and monomeric actin require the same sized gap between the filament barbed end and the boundary it pushes against, the Brownian Ratchet theory predicts that capping and elongation will respond to force in the same way. This means that, when the rate of filament elongation slows under load, the rate of capping also slows by the same amount. Filaments grow slower but they grow for a proportionally longer time and reach the same average length (Bieling et al., 2016). We directly tested a central prediction of the Brownian Ratchet model by creating a ‘bulky’ mutant capping protein that requires a larger-sized gap to bind the end of an actin filament. When we applied force to networks that contained the mutant capping protein, the rates of filament elongation and capping diverged sharply. This experiment illustrates the functional importance of the capping protein force response and provides the most direct experimental support for Brownian Ratchet models of force generation to date.

The matched force responses of filament elongation and capping suggest that the size of capping protein may be evolutionarily adaptive and subject to positive selection. A mismatch between the sizes of capping protein and actin would cause filaments to lengthen or shorten in response to force, both of which could be deleterious. Filament lengthening would impair force generation because longer filaments more easily bend and/or buckle, while filament shortening would weaken the network by decreasing filament density and network connectivity. In this way, a force-generating Brownian ratchet is controlled by a second, regulatory Brownian ratchet to form a balanced, self-assembly motor. Load-invariance of average filament length also means that this critical network parameter can be tuned independently by polymerases such as formins or Ena/VASP proteins or even WASP-family nucleation promoting factors.

## Supporting information

Supplemental Material for Li et al.

## Notes

### Competing Interest Statement

The authors have declared no competing interest.

