## Supplemental Material for Li et al. for "The molecular mechanism of load adaptation by branched actin networks"

### **Appendix: Brownian Ratchet Model of branched network assembly**

Based on the original Brownian Ratchet model for actin assembly (Peskin et al., 1993), the actin monomer association rate to filament free barbed ends growing against a load can be written a function of applied force as

$$(1)\quad R = \delta(\alpha e^{-\frac{f\cdot\delta}{K_B T}} - \beta)$$

, where δ is the Brownian-Ratchet gap size required for the addition of an actin monomer, α is the association rate in the absence of force , *f* is the force on an individual actin filaments, $K_B$ is the Boltzmann constant, *T* is the temperature in Kelvin and *β* is the dissociation rate. We assume that the addition of capping protein to the filament end is a very similar insertional process that, like monomer addition, requires an opening of a gap. The rate of capping should, therefore follow the same equation. For two differently sized capping proteins (wt CP and GST-CP) as in our experiment, we can relate their relative rate of incorporation (R) to their characteristic gap size (δ) via

$$(2)\quad \frac{R_{GST}}{R_{WT}} = \frac{\delta_{WT}}{\delta_{GST}} \cdot e^{\frac{-f(\delta_{WT}-\delta_{GST})}{K_B T}}$$

For filaments pushing against the load in orientations different from a normal (90°C) angle, the filament contact angle ($\theta$) has to be taken into account and the normalized ratio can be written as

$$(3)\quad e^{\frac{-f(\delta_{WT}-\delta_{GST})\cdot sin\theta}{K_B T}}$$

We can obtain the compressive force on an individual, growing filament via

$$(4)\quad f = \frac{F_{total}}{N_{load}^{BE}}$$

where $F_{total}$ is the total force generated by the number of free barbed ends sharing the load ($N_{load}^{BE}$). Within a branched network, the free barbed ends associating with the WH2 domain of the NPF are neither involved in elongation nor capping and thus do not contribute to active force generation. Therefore the average force on individual filaments has to be re-written as

$$(5)\quad f = \frac{F_{total}}{N_{total}^{BE} - N_{WH2}^{BE}}$$

where the $N^{BE}_{total}$ is the total number of free barbed ends and $N^{BE}_{WH2}$ is the number of free barbed ends associating with the WH2 domain of the NPF. Note that both of these quantities change when the network experiences external load (Figure 3, (Bieling et al., 2016)).

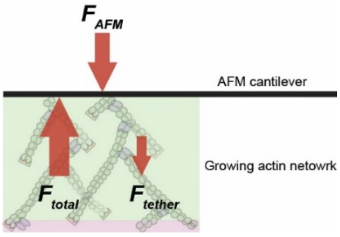


Finally, we assume that a filament end bound to the WH2 domain of the NPF exert a pulling or tether force ($f_{tether}$). This tethering force scales with the number of tethered ends and adds to the externally applied force via the AFM cantilever to resist movement. Therefore, the total load on the growing free barbed ends will be the sum of the total tethering force and applied AFM force:

$$(6)\quad F_{total} = F_{AFM} + F_{tether} = F_{AFM} + f_{tether} \cdot N^{BE}_{WH2},$$

Combining equations 3, 5 and 6, the normalized GST-CP to wt CP rate ratio becomes

$$(7)\quad Normalized\ rate\ ratio \sim e^{-\frac{F_{AFM}+f_{tether}\cdot N^{BE}_{WH2}}{K_BT\cdot(N^{BE}_{total}-N^{BE}_{WH2})}(\delta_{WT}-\delta_{GST})\cdot sin\theta}$$

All of these quantities, with the exception of the characteristic tethering force can be well estimated based on our measurements here and in (Bieling et al., 2016) as detailed in the following sections.

**Total number of free barbed ends ($N^{BE}_{total}$)**

To obtain the number of free barbed ends within the actin network ($N^{BE}_{total}$), we are using the previously measured actin polymerization rate per network area ($P_{actin}$ monomer/s/µm$^2$) and the measured network growth velocity ($V_{network}$ um/s) according to

$$(8)\quad V_{network} = \frac{P_{actin}\cdot A}{N^{BE}_{total}} \cdot \delta_{eff},$$

where $A$ is the network growing area and $\delta_{eff}$ is the effective ratchet size for one monomer association. For $P_{actin}$ we found a value of 7135 monomers/s/um$^2$ and $V_{network}$ is 5.69 um/min at 25 Pa (Bieling et al., 2016). The average contact angle between the filament ends and the load in the absence of external load can be assumed to be 54°, as a result of the characteristic angle of Arp2/3 branching. We therefore estimate $\delta_{eff} = 2.7\ nm \cdot Cos(\frac{72^o}{2})$. In our experiment setup, the network area is, 14x14 μm$^2$. Therefore, the total number of growing filaments (free barbed ends) of the whole network at 25 Pa can be calculated as 32211. The number of free barbed ends as a function of growth force was then obtained by scaling the measured relative density of free ends with this value (Fig. 1C, (Bieling et al., 2016)).

**Filament contact angle ($\theta$)**

The contact angle between actin filaments and the load changes in response to force to increase the actin density within the network (Bieling et al., 2016; Mueller et al., 2017). To calculate the filament contact angle, we are going to use the concept that the $N_{filament} \cdot Cos(\theta) \propto D_{actin}$, where $N_{filament}$ is the number of filaments and $D_{actin}$ is the actin density of the network. Also, in the growing actin network, the number of total filament is defined by the number of free barbed end ($N_{total}^{BE}$).

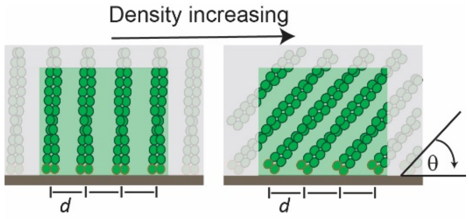


Therefore, the contact angle ($\theta$) can be written as

$$(9)\quad Sin(\theta) = Sin(\theta_0) \cdot normalized\left(\frac{D_{actin}}{N_{total}^{BE}}\right)$$

where the $\theta_0$ is the contact at zero force (54°). Both the $D_{actin}$ and $N_{total}^{BE}$ can be derived from data (Fig. 3D in (Bieling et al., 2016)).

**Number of free barbed ends associating with WH2 ($N_{WH2}^{BE}$)**

We show that the total WH2 occupied by the free barbed ends increases from 7.2% to 26.8% when the force increases from 0 to 1020 Pa (Figure 3). For simplicity, we assume a linear force dependence to calculate the number of attached ends according to

$$(10)\quad N_{WH2}^{BE} = \left(\left(\frac{26.8\%-7.2\%}{250-0}\right) \cdot F_{AFM} + 7.2\%\right) \cdot N_{total}^{WH2},$$

where $N_{total}^{WH2}$ is the total available WH2 on the surface.

**Tethering force of individual free barbed end-WH2 association ($f_{tether}$)**

The tethering force of free barbed end-WH2 association has been considered as external load-dependent in an exponential way [4]. For simplicity, we use single exponential decay in our model with

$$(11)\quad f_{tether} = f_{tether}^{0} \cdot e^{-aF_{AFM}}$$

where $f^0_{tether}$ is the tethering force at zero load and $a$ is the exponential decay constant.

The ratchet gap size for wild-type CP ($\delta_{WT}$) and GST mutant CP ($\delta_{GST}$) are 2.7 nm and 7 nm according to structural models (see Methods, (Funk et al., 2021; Kim et al., 2010; Narita et al., 2006). At this point, the $N^{WH2}_{total}$, $f^0_{tether}$, and $a$ are the free parameters in equation (7). The best fit of this model to our experimental data (see Figure 4F) yields $N^{WH2}_{total}{\sim}230000$, $f^0_{tether}{\sim}0.3\ pN$, and $a{\sim}0.025$. The first number is in a good agreement with previous data (Bieling et al., 2018). The tethering force has not been measured up to this point but 0.3 pN is in the range of weak protein-protein interactions; the exponential decay constant of this tethering force will need further quantitative investigation.

**Material and Methods:**

### 1. Protein biochemistry:

**1.1 Coverslip-immobilized proteins (NPFs and mCherry variants):** The coding sequence of Human WAVE1 lacking the N-terminal SH1 domain (AA 171-559, WAVE1ΔN) was codon-optimized for expression in E. Coli (GeneArt) and fused to an N-terminal mCherry-tag harboring an N-terminal Lys-Cys-Lys-(KCK-)tag (for surface immobilization) followed by a $His_{10}$-tag (for purification) and cloned into a modified pET vector containing a TEV-cleavable z-tag (REF). To prevent surface attachment via protein sites other than the N-terminal KCK-tag, endogenous Cysteine residues of WAVE1 (Cys 296 and 407) were replaced with Serine without affecting protein activity. A non-fluorescent version of this mCherry-NPF fusion construct was generated by introducing a Tyr71->Ser mutation in mCherry (darkCherry) to facilitate multicolor TIRF microscopy when direct visualization of the NPF was not necessary. For the detection of WH2 occupancy by FRET, a residue directly upstream of the $WH_2$ domain (Thr490 in WAVE1ΔN) was mutated to Cys and the N-terminal KCK- was substituted for a Sortase-($Gly_5$-)tag. For the purification of mCherry or dark mCherry lacking NPF activity, we introduced a STOP codon between the Cherry and NPF moiety. Proteins were expressed in E.Coli (Star pRARE) for 16h at 18°C and purified by IMAC over a HiTrap Chelatin column, followed overnight TEV cleavage on ice, ion-exchange chromatography over a Source Q (XK 16-20) column and gelfiltration over a HiLoad Superdex 200 column. The FRET NPF construct was labeled after ion-exchange chromatography with Alexa488-Maleimide at position 490 and then subjected to sortase-mediated peptide ligation of a Cys-containing peptide (CLPTEGG) to the N-terminal Sortase-($Gly_5$-)tag, followed by gelfiltration for the removal of free peptide. Proteins were SNAP-frozen in liquid nitrogen in storage buffer (20mM HEPES (pH=7.5), 150mM NaCl, 0.5mM TCEP, 0.1mM EDTA, 20% Glycerol).

**1.2 Actin:** Native, cytoplasmic actin from Amoeba castellanii was purified by ion-exchange chromatography and a cycle of polymerization-depolymerization as described previously (Hansen et al., 2013) and stored in filamentous form dialyzing against polymerization buffer (20mM Imidazole (pH=7.0), 50mM KCl, 1.5mM $MgCl_2$, 1mM EGTA, 0.5mM ATP, 0.5mM TCEP). 5ml fractions of the filamentous pool were depolymerized at a time by dialyzing into G-Buffer (2mM Tris-Cl (pH=8.0), 0.1mM $CaCl_2$, 0.2mM ATP, 0.5mM TCEP) for 1 week, followed by gelfiltration over a HiLoad Superdex 200 (XK16-60) column. Actin was kept in monomeric form after gelfiltration at 4C for up to two months. Actin was fluorescently labeled with Alexa488-Maleimide at Cys 374 as previously described (Hansen et al., 2013). For labeling actin with Atto540Q-NHS, the profilin-actin complex was formed in G-Buffer with a 1.5-fold excess of profilin. The complex was isolated by gelfitration over a Superdex 75 column in labeling buffer (2mM HEPES (pH=8.0), 0.1mM $CaCl_2$, 0.2mM ATP, 0.5mM TCEP), concentrated and labeled at reactive lysine residues by incubating with a 10-fold excess of the NHS-dye for 1h on ice. After quenching with Tris-Cl (2mM, pH=8.0), actin was polymerized by addition of 10x polymerization buffer and a small quantity (1% of total actin) of freshly sheared filaments. After polymerization for 1h at room temperature, filaments were pelleted by ultracentrifugation (20 min at 80 krpm in a TLA100.2 rotor) and then depolymerized in G-Buffer for 1 week in the dark. Depolymerized, labeled actin was then gelfiltered over a Superdex 75 column and stored on ice.

**1.3 Arp2/3 complex:** The native, bovine Arp2/3 complex was purified from calf thymus glands (PelFreez) by a series of ammonium sulfate precipitation and ion-exchange chromatography (DEAE, Source Q and Source S) steps followed by gelfiltration (Superdex

200) as described previously (Doolittle et al., 2013). Arp2/3 was fluorescently labeled by addition of 3-fold excess of maleide-dye conjugate, incubated for 1.5h on ice, quenched by adding DTT to 1mM and desalted into VCA Buffer A (5mM Tris-Cl (pH=8.0), 5mM NaCl, 1mM DTT, 0.2mM $MgCl_2$, 0.1mM ATP). To remove a small subfraction of Arp2/3, which irreversibly bound to the NPF after labeling, the complex was bound to a 5ml NPF affinity column (N-WASP VCA immobilized on a HiTrapNHS resin) and eluted by a 10CV gradient to VCA Buffer B (5mM Tris-Cl (pH=8.0), 5mM NaCl, 0.2mM $MgCl_2$, 0.1mM TCEP, 0.1mM ATP). Peak fractions were pooled, concentrated and gelfiltered over a Superose 6 column. Proteins were SNAP-frozen in liquid nitrogen in storage buffer (5mM HEPES (pH=7.5), 50mM NaCl, 0.5mM $MgCl_2$, 0.5mM TCEP, 0.5mM EGTA, 0.1mM ATP, 20% Glycerol).

**1.4 Capping protein:** To generate wt CP, the α1 and β2 isoforms of murine heterodimeric capping protein were cloned into pETM20 and pETM33, respectively. To generate fluorescently tagged, wt CP, an N-terminal SNAP-tag (Keppler et al., 2003) was fused to the beta subunit. To construct larger sized CP dimers ("bulky mutants") we fused either a GST-tag (aa 1-217 of Glutathione S-transferase from Schistosoma japonicum) to the N-termini of both CP subunits or a NusA-tag (full length from Escherichia Coli) to the alpha subunit. GST and CP domains were separated by a $(Gly)_5$Thr- (21.5 Å) spacer, whereas NusA and CP domains were separated by a GlyThr-(7.2 Å) spacer. GST-CP αwas cloned into pETM20, whereas GST-CPβ and NusA-CPα were cloned into pETM11. To generate fluorescently tagged GST-CP, an N-terminal SNAP-tag was additionally fused to the GST- CP βchimera. To generate fluorescently tagged NusA-CP, an N-terminal SNAP-tag was additionally fused to the NusA-CPα chimera. Proteins were co-expressed in corresponding pairs of alpha and beta subunit combinations in E.Coli (Rosetta) for 16h at 18C and purified by IMAC over a 5ml HiTrap Chelating column followed by overnight TEV/Prescission cleavage of the N-terminal His-tags on ice. After desalting over a HiLoad Desalting column, uncleaved protein and free tags were removed by recirculation over the IMAC column. The flow through was subjected to ion-exchange chromatography over a Mono Q column and gelfiltration over a Superose 6 column. Proteins were SNAP-frozen in liquid nitrogen in storage buffer (10mM Tris-Cl (pH=7.5), 50mM NaCl, 0.5mM TCEP, 20% Glycerol). Addition of the N-terminal tags (SNAP-, GST-, NusA- or combination of those) did not affect capping activity in the absence of force as measured by polymerization of pyrene-actin in bulk (Supplemental Figure 3) or capping in single filament TIRFM assays.

**1.5 Profilin:** Human profilin 1 was expressed and purified as previously described (Bieling et al., 2016) and SNAP-frozen in liquid nitrogen in storage buffer (10mM Tris (pH=8.0), 50mM KCl, 1mM EDTA, 0.5mM TCEP, 20% Glycerol)

**1.6 Ezrin-ABD:** The C-terminus of human Ezrin (aa 553-586) followed by a 13aa Gly-rich linker and a C-terminal KCK-motif was cloned into pGEX-6P-2, expressed in E.Coli (Rosetta) for 8h at 25C, purified over a GST Trap column followed by overnight GST-Prescission cleavage on ice and desalting. Desalted protein was filtered over a GST Trap column to remove free GST and GST-Prescission. The flow through was gelfiltered over a Superdex 75 column and SNAP frozen in liquid nitrogen in in storage buffer (10mM Tris (pH=8.0), 150mM KCl, 0.5mM TCEP, 20% Glycerol)

**1.7 Myotrophin/VI:** Full length, human myotrophin was cloned into a modified pETM11 vector containing a TEV-cleavable, N-terminal $His_{10}$-tag and expressed in expressed in E.Coli (Rosetta) for 8h at 25C, purified over a HiTrap Chelatin column followed by overnight TEV cleavage on ice and desalting. Desalted protein was filtered over a HiTrap Chelatin column to remove free His-tag and TEV. The flow through was gelfiltered over a Superdex 200 column and SNAP frozen in liquid nitrogen in in storage buffer (20mM HEPES (pH=7.5), 150mM KCl, 0.5mM TCEP, 20% Glycerol)

**2. Surface functionalization and protein immobilization:**

**2.1 Coverslip functionalization, photolithography and protein immobilization:** Glass coverslips (22x22 mm, #1.5, high precision, Zeiss) were functionalized and patterned as described previously (Bieling et al., 2016). Briefly, surfaces were rigorously cleaned by consecutive incubation in 3M NaOH and Piranha solution (3:2 concentrated sulfuric acid to 30% hydrogen peroxide) followed by silanization with (3-Glycidyloxypropyl)trimethoxysilane. Silanized surfaces were then passivated by reacting with diamino-PEG. Subsequently, exposed amino groups were reacted with a heterobifunctional crosslinker (BMPS) to create PEG-maleimide coated coverslips, which were subjected to UV-microlithography using a chrome-on-quartz photomasks, which selectively protected maleimide groups within chrome-covered areas from UV exposure. Micropatterned PEG-maleimde coverslips were then loosely attached to flow chambers constructed of PLL-PEG passivated microscopy counter slides and thin PDMS stripes (flow cell volume=40μl). For the immobilization of NPF on micropatterned PEG-maleimide coverslips, protein aliquots of KCK-Cherry- WAVE1ΔN (NPF) and KCK-Cherry (mock protein) were rapidly thawed and und pre-reduced with 1mM beta-mercaptoethanol for 30min on ice and then desalted twice into Immobilization Buffer (20mM HEPES (pH=7.5), 300mM NaCl, 0.5mM EDTA). Protein concentration was determined by $Au_{280nm}$ and NPF protein mix was prepared by diluting desalted proteins to 10mM total in immobilization buffer, followed by direct incubation for 25min at room temperature with the freshly patterned PEG-maleimide coverslip in the flow cell contained in a humidified chamber. The NPF density was controlled by adjusting the relative percentage of KCK-darkCherry-WAVE1ΔN (NPF) and KCK- darkCherry (mock protein). Coverslips were prepared using a percentage of 60% NPF and 40% mock protein. For the FRET experiments determining the WH2 occupancy, coverslips were prepared using 50% KCK-darkCherry-WAVE1ΔN, 10% CLPTE-darkCherry-WAVE1ΔN (Alexa488-Cys490) and 40% KCK-darkCherry. After protein immobilization, flow cells were washed with 6 flow cell volumes wash buffer (20mM HEPES (pH=7.5), 300mM NaCl, 0.5mM EDTA, 5mM beta-mercaptoethanol), incubated for 3min to quench residual maleimide groups, washed with 6 flow cell volumes storage buffer (20mM HEPES (pH=7.5), 300mM NaCl, 0.5mM EDTA, 2mM TCEP) and stored at 4°C in a humid container for up to 5 days.

**2.2 Cantilever functionalization:** Ezrin-coated AFM cantilevers were prepared as described previously (Bieling et al., 2016). Tipless, uncoated cantilevers were chemically cleaned by incubating in Piranha solution (3:2 concentrated sulfuric acid to 30% hydrogen peroxide), washed, transferred to custom-built PDMS incubation chambers and functionalized by incubating for 1.5 h in Silane-PEG5000-Maleide (Nanocs, freshly resuspended to 2% (w/w) in 95% ethanol, 5% water, pH=5.0) at room temperature. The cantilevers were then washed twice in excess ethanol, dried for 1h at 75°C and washed with ultrapure water. Ezrin-ABD was diluted to 20uM in cantilever buffer (2mM Tris-Cl, pH=8.0) and immobilized on PEG-Maleimide-functionalized cantilevers by overnight incubation at 4C in custom-built PDMS incubation chambers. Immediately before the experiment, Ezrin-coated cantilevers were washed in excess cantilever buffers and dried.

**3. Fluorescence and atomic force microscopy system:**

**3.1 TIRFM-AFM system:** Imaging was performed on an Observer.Z1 (Zeiss) microscope equipped with a total internal reflection fluorescence (TIRF) slider (Zeiss), a TIRF objective (PlanApochromat 100X 1.46 TIRFM, Zeiss) and a cooled charge-coupled device camera (iXon888, Andor). Fluorescence excitation was accomplished by three diode-pumped solid-state laser lines (488, 561 and 644nm), which were controlled using an acousto-optical filter and coupled into a single fiberoptic light guide (custom laser launch, Spectral Applied Research). Micro-Manager (Edelstein et al., 2010) was used to control the shutters, acousto-optical filter, dichroic mirrors and camera. Laser intensity and exposure was minimized to avoid photo-bleaching. For bulk multi-color fluorescence measurements of dendritic network component densities, images (300ms exposure time) were taken at custom intervals of increasing time (5-30s to avoid bleaching in networks growing with reduced velocity at elevated forces). Fast, one-color imaging of single molecules was performed at an increased frame rate of 10 frames/s and a 100 ms exposure time (''streaming'' mode).

Force measurements were performed using commercial AFM system (BioScope Catalyst, Bruker) modified as described in detail in (Bieling et al., 2016). Briefly we 1) replaced the original AFM photodiode detector with a position sensitive device (PSD, Pacific Silicon Sensor, DL100-7PCBA3) to obtain a larger dynamic force range, 2) constructed a custom cantilever holder with low cantilever angle (~3$^o$) to closely match the idealized geometry of two parallel planes and also to prevent slippage between the AFM cantilever and actin network, 3) constructed a custom sample holder that could prevent evaporation, and 4) utilized a customized setup to perform micro-rheology measurements (Alcaraz et al., 2003; Mahaffy et al., 2000). All the AFM electronic signals from PSD were pre-processed by an electronic filter (Krohn-Hite, 3362) set to dc low-pass at 30 Hz. Custom-written software in LabView was used for signal processing, data acquisition, and piezo stage control.

### 4. Denditic network assembly assays:

**4.1 General dendritic network assembly assay:** Flow cells of micropatterned, NPF-coated coverslips were washed with twice with 250ul of ultra-pure water (Milli-Q grade) and disassembled by removal of the coverslips. Excess water was removed by a brief (5s) spin on a spin coater. Drying did not affect NPF activity if the coverslip was not kept in air for >30min. The coverslip was fixated on a custom-built sample holder by adhering to a thin PDMS O-ring and the whole assembly was transferred to the microscope stage. An ezrin-coated AFM cantilever was immobilized with a drop of hot paraffin wax on a custom built cantilever holder and then attached to the AFM head, which was mounted on the microscope and lowered to close proximity to the coverslip.  100 ml assembly buffer (20mM HEPES (pH=7.0), 100mM KCl, 20mM beta-Mercaptoethanol, 1.5mM $MgCl_2$ , 1mM EGTA, 1mM ATP, 0.5mg/ml beta-casein, 10nM Alexa488-labelled actin) were added in between coverslip and cantilever holder. Low amounts of labelled actin were included in the buffer to visualize the NPF patterns indirectly via the binding of actin monomers. 80ul of mineral oil containing 20mg/ml Cithrol DPHS (to passivate the oil-buffer interface) was overlaid onto the buffer to seal if from air exposure. The Optical Lever Sensitivity (OLS) is characterized by measuring the force-distance curve in contact with the hard glass surface, prior to every measurement. An NPF pattern was then positioned at an axial distance of 3um directly under the AFM cantilever via the motorized stage and the x- and y-piezoelectric stage control. Actin network growth was finally initiated by addition of 50ul of network proteins in assembly buffer (final concentration: 5uM actin, 5uM profilin, 100nM Arp2/3, 100nM CP if not indicated otherwise). Synchronously, multicolor TIRFM time-lapse imaging was initiated. For bulk fluorescence, multicolor TIRFM experiments, the protein mix was supplemented with

1%Alexa488-actin, 5% Alexa647-Arp2/3 and 15% TMR-SNAP-CP. After the height of the growing network reached the cantilever (as indicated by cantilever displacement and a rise in force), the force was kept constant at a defined setpoint by engaging the force-feedback mechanism ("force-clamp mode"). The force was maintained until both network fluorescence and growth velocity reached a steady state, upon which the force was changed to a higher setpoint. This cycle was repeated until network growth was slowed to velocities <200 nm/min, close to mechanical stall. Network growth did not exhibit hysteresis, hence the order or duration by which the individual forces were applied did not affect the growth velocity.

**4.2 Single molecule dendritic network assembly assay for TIRFM-AFM:** Assays were carried out as described in the section 4.1 with the following exceptions: For single color, single molecule imaging, assembly buffer was supplemented with an oxygen scavenger system (40mM glucose, 125 ug/ml glucose oxidase, 40mg/ml catalase) and 2mM Trolox and the protein mix contained 0.02% (1 in 5000) Alexa647-Arp2/3.

**4.3 FRET assay for the determination of the WH2 occupancy of the NPF:** Assays were carried out described in section 4.1 with the following exceptions: Coverslips were functionalized using a low percentage of the NPF FRET construct (CLPTE-darkCherry-WAVE1 ΔN(Alexa488-Cys490), see section 2.1). Reactions were scaled down to 100ul volume (in comparison to 150ul for standard conditions). Assembly buffer was supplemented with an oxygen scavenger system (40mM glucose, 125 ug/ml glucose oxidase, 40mg/ml catalase), 2mM Trolox and the protein mix contained only Alexa647-Actin (1% of total) as fluorescent label. The load was maintained at 1020 Pa until the network reached steady state growth and a minimum height of >3um. Network growth was then arrested and capping was inhibited by carefully diluting the reaction (100 ul total) by adding 200ul fixing buffer (assembly buffer containing Latrunculin B (15uM final), Phalloidin (15uM final), Myotrophin (5uM final, competitive, high-affinity CP inhibitor (Bhattacharya et al., 2006))), profilin (7.5uM final) and Atto540Q-actin (7.5uM final). The fraction of quencher-labeled actin (Atto540Q-actin) after arrest was thus 7.5uM of 10uM total. For control experiments in the absence of a dendritic network, the assembly buffer was supplemented with 15uM Latrunculin B before the experiment to prevent actin polymerization.

**4.4 "Spike-in" experiments using larger sized capping protein variants ("bulky mutants"):** Experiments were carried out as described in section 4.1 with the following exceptions: The overall CP pool (100nM) consisted of 90% unlabeled, wt CP, 5% TMR-SNAP-CP and either 5% Alexa 647-SNAP-GST-CP or 5% Alexa 647-SNAP-NusA-CP. This limited the influence of the lowered capping rate of the size mutant at elevated forces on the overall network assembly kinetics.

### 5. Data analysis:

**5.1 Quantification of network growth velocity:** Constant growth forces were applied to a growing network under AFM force clamp control. The growth velocity of the network was determined by the slope of height-time curve at individual constant growth forces. However after switching to a new growth force, the network needed time to adapt the new growth force to reach constant growth. Therefore, the slope is not considered for growth velocity until the network reached the steady constant growth where the slope is constant.

**5.2 Quantification of bulk fluorescence intensities from TIRFM and confocal imaging:** The mean intensities of all network components (actin, Arp2/3, CP) from multicolor, time-lapse TIRFM images were quantified via ImageJ (ROI Manager->Multi Measure function) from square region of interests (ROIs) matching the network area. Background intensity was determined from adjacent regions (10um distance) of the same size and subtracted from the network intensity. For Arp2/3 and actin, a small (<30% of total intensity in the absence of force for TIRFM imaging, <5% for confocal imaging) fraction of fluorescence in the network area is due to binding to the NPF in addition to the actin network. The intensity of this signal was quantified during the initial lag phase preceding actin network nucleation and subtracted from the network intensity. The fluorescence intensities were plotted as a function of time together with the height of the sample as well the counterforce. Mean fluorescence intensities at were then calculated by averaging over the fluorescence signal during steady growth at a constant force. The variance in fluorescence intensity during these steady state periods was very low (SD<2% total).

**5.3 Single molecule tracking and classification:** For detection and tracking of single Arp2/3 molecules in TIRF time-lapse images, we used the u-track software package (Jaqaman et al., 2008). After complete tracking, an additional step classifies tracks into productive (molecules that are incorporated into the network and continuously grow out of the TIRF microscopy field of view as indicated by a progressive drop in fluorescence intensity) or unproductive (stuck and/or blinking molecules at constant intensity and position). This is done in a semi-automated process: All individual tracks of minimum length 5 frames (= 500 ms) are randomly distributed amongst six biological experts. Each expert subsequently classifies all individual tracks of his share as productive or unproductive. In this step, 10% of all tracks are classified by two experts independently to estimate the classification uncertainty. This led to a fraction of tracks of at least 90% over all sample clips that is associated to the same class by both experts. In order to efficiently process significant amounts of microscopy data, we further automated our analysis procedure, by calculating a set of 10 feature parameters for each track. All features are based on the dynamics of intensity and position of each track. The set of features in combination with the combined classification results of the experts was used subsequently for training a supervised random forest classifier. Cross validation yielded correct classification in at least 82% of all tracks. In order to further improve this performance, we followed an active learning strategy in which borderline cases (i.e. tracks for which the decision trees in the forest do not agree well in their classification decision) are decided by an expert. Using cross validation of this semi-automated procedure, the amount of manually classified tracks is tuned to yield a comparably high correct classification performance as the group of experts (i.e. > ~90%). After classification, productive tracks are used in further analyses exclusively.

**5.4 Determination of bulk nucleation rates from single molecule calibration experiments:** The mean event rates of productive network incorporation (in counts per network per second) was determined for Arp2/3 from single molecule “spike-in” experiments at 25 Pa and multiplied by the respective labeling ratio to yield the total nucleation rate at this load.

**5.5 Quantification of the WH2 occupancy of the NPF by FRET TIRFM imaging:** The fluorescence of donor-labeled NPF (CLPTE-darkCherry-WAVE1ΔN(Alexa488-Cys490)) was plotted as function of time after addition of quencher-labeled actin (Atto540Q-actin). The data was fitted to a sum of two exponential functions:  with the rapid phase caused by FRET (binding of quencher-labeled actin to the WH2 domain of the NPF) and the slow phase attributed to bleaching of the donor. The bleaching rate ($k_2$) was independently determined in control experiments and fixed when fitting the FRET data. The amplitude of the fast, FRET phase ($I_{FRET}$) was determined for three cases: a) In the absence of a dendritic network, b) in the presence of a dendritic actin network that was assembled in the absence of force (0 Pa) or c) in the presence on a dendritic actin network that was assembled at a defined load force of 1020 Pa. Assuming that all WH2 domains are free to interact with monomers in the absence of a dendritic network (a), we calculated the amount of blocked, occupied WH2 domains in the other cases (b and c) by the relative decrease in $I_{FRET}$.

### Supplemental Figures:

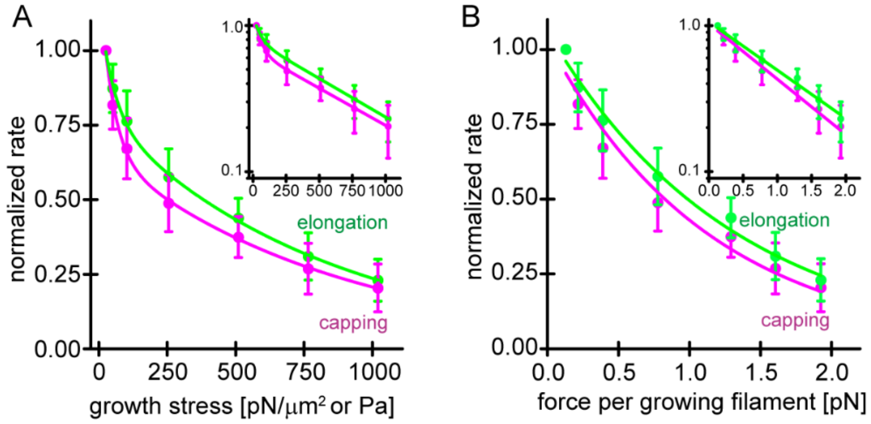


**Supplemental Figure 1: Load dependence of filament elongation and capping**

**A)** Average rates of filament elongation (green) and capping (magenta) as a function of total load on the network as in Figure 1D. Lines are fits to double exponential decay functions. Inset: Semi-logarithmic plots of the same data. **B)** Average rates of filament elongation (green) and capping (magenta) as a function of force per growing filament end. Lines are fits to single exponential decay functions. Inset: Semi-logarithmic plots of the same data. Error bars are SEM.

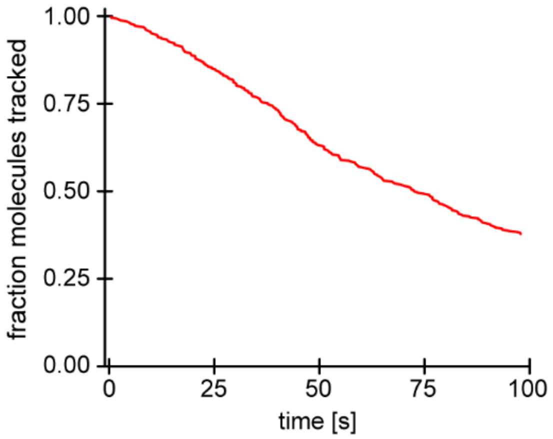


**Supplemental Figure 2: Tracking and bleaching control for single molecule imaging of Arp2/3 nucleation.**

Survival function of tracked molecules for surface-immobilized Arp2/3 complexes as a function of time under the same imaging conditions as for network-associated Arp2/3. Loss of molecules is based on both bleaching and loss of tracking.

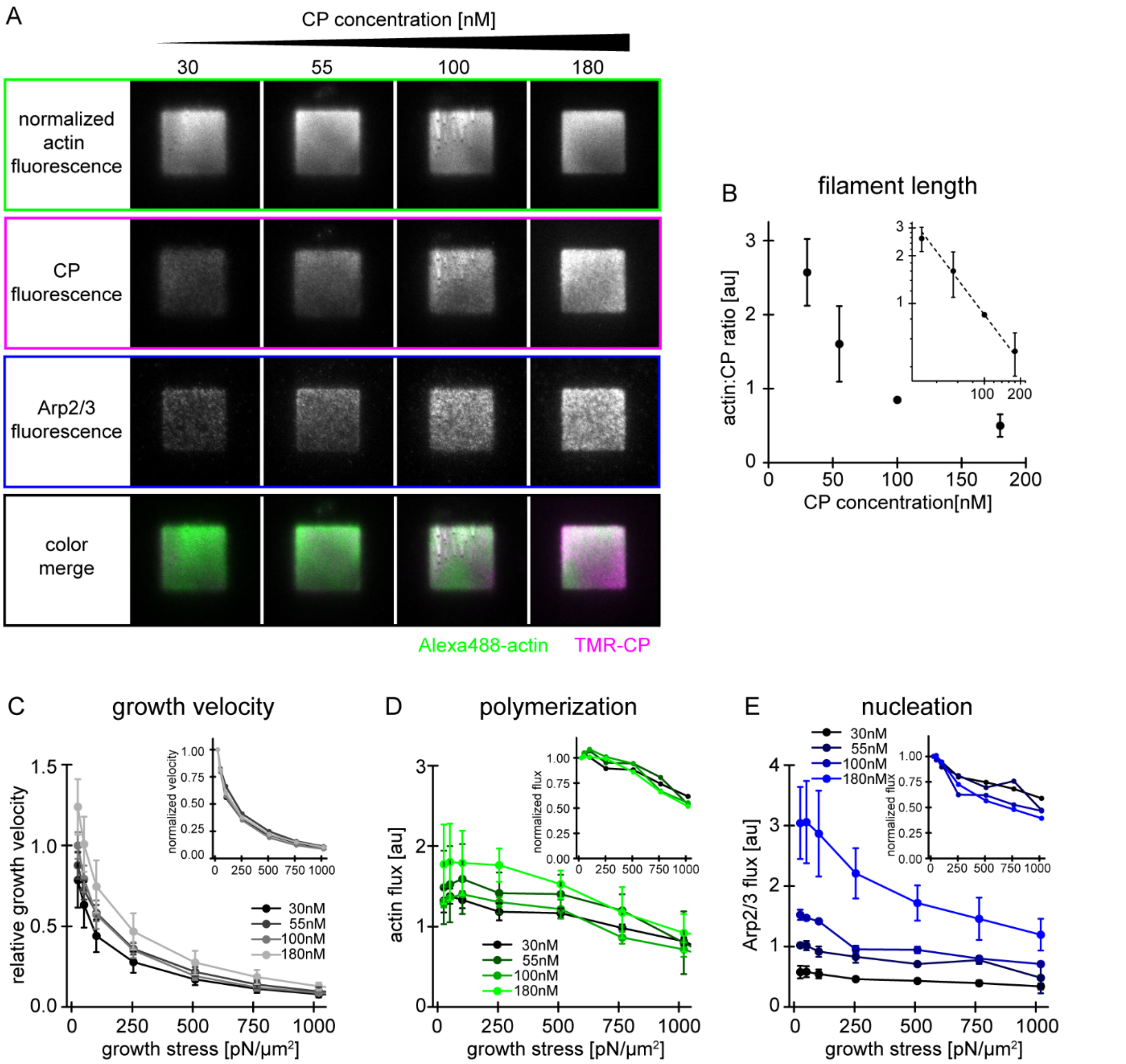


**Supplemental Figure 3: Characterization of network assembly at various CP concentrations. A)** Representative TIRFM images of individual protein constituents as indicated in growing actin networks loaded with low (25 Pa) growth stress. Conditions were 5uM profilin-actin (1%Alexa488-actin), 100nM Arp2/3 (5% Alexa647-Arp2/3) and CP concentrations as indicated (15% TMR-CP). **B)** Relative filament length as calculated from the actin:CP ratio as a function of the total CP concentration. Inset: Double-logarithmic plot. **C)** Relative network growth velocities normalized to the standard condition (25Pa growth stress at 100nM CP) for networks growing in the presence of various CP concentrations as indicated. Inset: same data but normalized to the growth velocity at 25Pa at the same CP concentration. **D)** Relative actin flux determined by the product of the bulk Alexa488-actin fluorescence intensity (as in A) and the corresponding growth velocity (as in C). Inset: Same data but normalized to the actin flux at 25Pa at the same CP concentration. **E)** Relative Arp2/3 flux determined by the product of the bulk Alexa647-Arp2/3 fluorescence intensity (as in A)

and the corresponding growth velocity (as in C). Inset: Same data but normalized to the Arp2/3 flux at 25Pa at the same CP concentration. Error bars are SD.

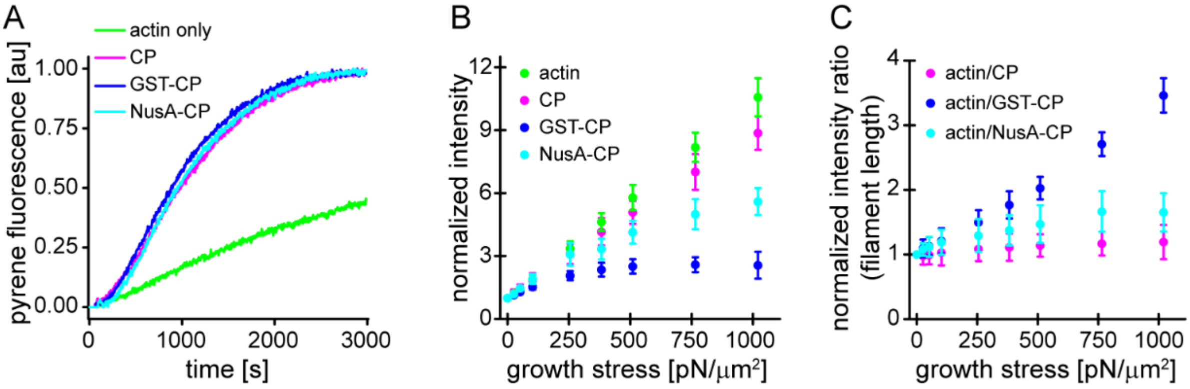


**Supplemental Figure 4: Characterization of CP variants.** A) CP (wt, magenta), GST-CP (bulky variant 1, blue) and NusA-CP (bulky variant 2, cyan), similarly stimulate the generation of free pointed ends of actin (2μM containing 5% pyrenyl-actin) in the absence of profilin. Green trace is the negative control containing only actin. CP concentration was 30nM total for all cases. B) Mean Alexa 488-actin, TMR-CP(wt, magenta), Alexa 647-GST-CP (bulky variant 1, blue) or Alexa 647-NusA-CP (bulky variant 2, cyan) intensity normalized to the intensity of an adjacent unloaded network as a function of load. C) Mean fluorescence intensity ratios of actin/CP(wt) (magenta), actin/GST-CP(bulky variant 1, blue) or actin/GST-CP(bulky variant 2, cyan) normalized to the intensity ratio of an adjacent unloaded network as a function of load. Error bars are SD.
